# Cross-validating the electrophysiological markers of early face categorization

**DOI:** 10.1101/2024.07.03.601129

**Authors:** Fazilet Zeynep Yildirim-Keles, Lisa Stacchi, Roberto Caldara

## Abstract

Human face categorization has been widely studied using electroencephalogram (EEG) and event related potentials (ERP). Within this context, the N170 ERP component has emerged as the earliest and most robust neural marker of face categorization, documented in thousands of studies conducted over nearly three decades. However, in the last decade there has been a surge in research using the fast periodic visual stimulation (FPVS) methodology to investigate face categorization. FPVS studies have consistently reported robust bilateral face categorization responses over the occipitotemporal cortex with a right hemispheric dominance, closely mirroring the N170 scalp topography. Yet, the question remains whether the neural response elicited in FPVS can be considered a proxy for the N170 or if it might be driven by different components. To address this issue, we recorded the electrophysiological signals of human observers who viewed natural images of faces and non-face objects during FPVS and three different ERP paradigms. We quantified the FPVS response in the frequency domain and extracted ERP visual components, including the P1, N170 and P2 in response to face stimuli, from both the ERP paradigms as well as the time domain of the FPVS response. Our results revealed little relationship between any single ERP component and the FPVS frequency response. Across methodologies, only the peak-to-peak differences between N170 and P2 components significantly and consistently explained the FPVS frequency response. Our data show that the FPVS frequency response is not singularly contingent on any isolated ERP component, such as the N170, but rather reflects a later complex neural integration. These findings raise crucial methodological and theoretical considerations on the relationship between FPVS and ERP responses, urging caution when interpreting the neurofunctional role of both electrophysiological signals. While both markers are indicative of human face categorization, they appear to capture different stages in this cognitive process.

## 1. Introduction

How the human brain achieves visual object categorization is a fundamental question in cognitive neuroscience. In this framework, human faces hold a distinct significance, as their efficient categorization represents an extraordinary and biologically relevant feat for the visual system that has likely been subject to evolutionary pressure. As a visual input, faces represent a high-dimensional stimulus comprised of various features located within a specific spatial configuration including subtle differences in shape, texture, and colour. The effective processing of faces is critical as they stand out as the most familiar and socially significant visual cue in our visual environment. For successful face versus non-face visual categorization, faces should be discriminated from objects and then generalized across other face exemplars. Discriminating faces from non-face stimuli is a non-trivial task for the visual system because of the diverse range of stimuli in the natural world that exhibit visual features or properties reminiscent of faces, such as certain man-made objects like cars or buildings. Generalizing across various face exemplars is also a challenging task, considering the human visual system encounters numerous individual face instances, each presenting a broad spectrum of physical diversity, such as variations in relative size, gender, ethnicity, emotional expression, and age.

To gain deeper insight into how the brain achieve this remarkable feat of face categorization, several neuroimaging techniques have been used over the decades. A very popular technique is the electroencephalogram (EEG), which thanks to its non-invasive and high temporal resolution has been widely used to explore the unfolding of face categorization (and in general, face processing). Specifically, in the context of EEG, event-related potentials (ERPs) elicited by transient stimulation have proven to be highly useful to understand how face categorization is impacted by intrinsic (e.g., expectation) as well as extrinsic (e.g., light, head orientation) factors. The N170 ERP is the most widely investigated neurofunctional marker of early face categorization, with more than a thousand studies over nearly 30 years (Bentin et al., 1996; Bötzel et al., 1995; George et al., 1996). The N170 is a negative deflection occurring between 130-200 ms on the bilateral occipitotemporal electrodes that is larger in response to faces compared to other types of visual stimuli, such as objects, words, or scrambled faces. Even though the interpretation of the N170 component and its specific links to different aspects of face perception are still debated, there is a consensus that this component is a valid electrophysiological marker of face-sensitivity (for a review see: Rossion & Jacques, 2011).

Despite their usefulness in studying the unfolding of face processing, ERPs are not immune to challenges. ERPs have low signal-to-noise ratio (SNR), necessitating the collection of numerous trials for reliable measurement. Additionally, comparisons between conditions (e.g., face vs non-face stimuli) typically necessitate post hoc analyses, involving the subtraction of one response from another. This approach assumes that the two signals were acquired under identical circumstances, which is often not the case. Variability in attention levels, fatigue, or other external factors can introduce confounds, challenging the validity of these comparisons. Lastly, the extraction of ERPs can be influenced by subjective decisions made by experimenters. For instance, choices regarding whether to extract the mean peak or the maximum peak, or to base the selection of peak latency on the average across electrodes or the activity of the strongest electrode, can introduce variability into the analysis, potentially impacting the reliability and interpretability of findings.

An emerging alternative method that circumvents all these issues is a paradigm named oddball fast periodic visual stimulation (FPVS; Retter & Rossion, 2016; Rossion, 2014; Rossion et al., 2015), which relies on the ability of the brain to synchronize to a periodic stimulation producing steady state visual evoked potentials (SSVEP) (Adrian & Matthews, 1934; Regan, 1966). Within the context of face perception, this approach has been adapted to provide a direct investigation of different types of face processing, bypassing the need for post hoc contrasts between conditions and has led to hundreds of publications over the recent years. The oddball FPVS method involves presenting a series of stimuli (i.e., base) at a constant frequency rate, with periodically intervening stimuli (i.e., oddball) that differ from the base stream along a specific dimension (e.g., identity, expression, category). For example, to investigate face categorization, the base stimuli consist of images of non-face objects, while the oddball stimuli consist of images of faces. This method produces distinct SSVEP responses at two different frequencies, a response at the base frequency and another response at the oddball frequency. Face categorization is then quantified by the amplitude of the oddball frequency. The oddball FPVS paradigm operates on the principle that if the visual system detects a difference between the base and oddball stimuli, it will elicit different responses to each. Compared to the ERP method, oddball FPVS allows for a direct investigation of the process of interest, as the measured response is a differential response between the base and oddball images. Additionally, the SSVEP response has high SNR, as it is located at a narrow frequency band while background noise is spread over multiple frequency bands. Furthermore, as the frequency of the response is directly determined by the experimenters’ choice of stimulation rate, its quantification can be performed unambiguously, bypassing the need of subjective choices. Altogether, these characteristics make the oddball FPVS paradigm a highly promising and versatile tool in various contexts ranging from testing healthy populations to assessing clinical patients and infants.

Despite more than a decade of numerous studies employing the oddball FPVS paradigm, the exact nature of what the FPVS paradigm and its SSVEP responses are capturing remains elusive. Within the context of neural facial identity discrimination (FID), the oddball response was suggested to be related to the well-known N170 ERP component (Rossion et al., 2012). Specifically, it has been shown that the SSVEP-FID response is reduced for inverted compared to upright faces (Rossion et al., 2012), an effect that has also been reported for the adaptation effect on N170 (Jacques et al., 2007). Additionally, the same study found that the phase of the response in the frequency domain and the latency of the N170 component extracted in the time domain were comparably modulated by this experimental manipulation (Rossion et al., 2012). Furthermore, the SSVEP-FID response exhibits a scalp topography highly resembling to the one elicited by the face-sensitive N170 component, with stronger neural responses over the bilateral occipitotemporal electrodes (Liu-Shuang et al., 2014; Rossion et al., 2012).

In the context of face categorization (i.e., when base stimuli are objects and oddball stimuli are face) such *direct* comparisons between the ERP N170 and SSVEP frequency responses have not been conducted yet. Despite the numerous electrophysiological studies using one or the other approach, whether the N170 neural response are tapping into the same neural network of the SSVEP responses is unknown. Nevertheless, studies have explored the temporal domain of face-categorization responses in oddball FPVS paradigm. This revealed the presence of a complex waveform consisting of peaks and through that resembles, in topography and latency, the traditional P1, N170 and P2 ERP components (Hauk et al., 2021; Quek & Rossion, 2017; Retter & Rossion, 2016; Rossion et al., 2015). In addition to the well-established N170 ERP component, the P1 and P2 ERP components have also been frequently observed in studies of face processing. The P1 component, a positive peak typically occurring around 100 milliseconds after stimulus onset, reflects early visual processing (Herrmann et al., 2005; Jemel et al., 2003), while the P2 component, a positive peak following the early face-sensitive N170, is thought to be involved in later stages of face processing (Latinus & Taylor, 2005, 2006). Whether the SSVEP frequency response is related to any of these peaks that can be observed in its time domain remains an open question. Similarly, it remains unknown whether the SSVEP response is associated with any transient ERP component recorded using traditional ERP paradigms, where a face stimulus is presented in isolation rather than within a sequence of images.

In this study we aimed bridge this important theoretical and methodological gap in our understanding of both neural face-sensitive categorization responses, by empirically and *directly* relating SSVEP response to its time domain and to transient ERP responses. To this aim, we collected EEG data from human participants while they viewed natural images of objects and faces. In the oddball FPVS paradigm, participants saw images of non-face objects, which were presented at a high constant frequency rate. Within the stream of base images, oddball face stimuli were periodically introduced every fifth image. Face categorization was quantified in the frequency domain as the strength of the EEG response at the oddball frequency. Additionally, we extracted ERPs from the time domain of the signal in correspondence to the onset of face stimuli. Transient ERPs were recorded using two different paradigms. In the first approach we followed the traditional paradigm of presenting images of faces and objects one at a time and extracted the *isolated* face responses (i.e., ERPs elicited by isolated faces). However, the relationship between SSVEPs and transient ERPs might not be immediately apparent as these two responses are the results of different processes. As described by Rossion (2014), the SSVEP is a differential neural response obtained by contrasting non-face and face objects. This results in a face-specific response. On the other hand, the traditional practice of presenting faces which are temporally and spatially isolated from other stimuli, triggers a face response that is potentially contaminated by other processes. Therefore, we had two additional ERPs measures. First, we obtained *differential* face responses by subtracting the ERPs elicited by isolated objects from the ERPs elicited by isolated faces. Second, to bring the ERPs paradigm closer to the FPVS conditions, we also examined the response triggered by a modified ERPs paradigm. In this case, face stimuli were not presented in complete isolation, but were preceded by a short sequence of non-face images (i.e., *contextual* face responses). We reasoned that this should trigger ERPs that would more closely relate to the SSVEP response as in both cases the response to face-stimuli would be a differential wave.

Across the four time-domain data (i.e., FPVS time domain, isolated ERPs, differential ERPs, and contextual ERPs), we explored the relationship between SSVEP-frequency response and three main peaks corresponding to the P1, N170 and P2. We focused on these components as they are the most prominent peaks that can be observed in the SSVEP-time response and might therefore be the major contributors to the response in the frequency domain. Additionally, we also investigated the peak-to-peak difference between P1 and N170 and between N170 and P2. Our reasoning is based on how the frequency domain is extracted from the SSVEP response. Specifically, the signal in the time domain undergoes a Fast Fourier Transforms that determines the presence of sinusoidal wave of different frequencies in the signal. As sinusoidal waves are characterized by peaks and through, it is reasonable to expect that the amplitude in the frequency domain is determined not only by one maximum or minimum in the signal but by a combination of them. Therefore, only considering baseline-to-peaks might provide only partial answers. Finally, we explored whether these neural responses would relate to behavioral performance at a test of face recognition, the CFMT+ (Russell et al., 2009). This investigation further elucidates the practical implications of our findings, offering insights into the potential applications of SSVEP responses in real-world contexts.

To anticipate our findings, our data revealed little relationship between any single ERP component and the FPVS frequency response. Across methodologies, only the peak-to-peak differences between N170 and P2 components significantly and consistently explained the FPVS frequency response. Our data show that the FPVS frequency response is not singularly contingent on any isolated ERP component, such as the N170, but rather reflects a later complex neural integration.

## 2. Methods

### 2.1 Overview

We recorded the electrophysiological signals of human observers who viewed natural images of objects and faces in ERP and FPVS paradigms. Then, from the same observers we recorded behavioural responses to a facial identity recognition task. We assessed how the FPVS frequency response relates to its own time-domain components, as well as components extracted from three ERP measures including *isolated* faces in which faces were presented in isolation, *differential* faces in which isolated non-face object images were subtracted from the isolated face images, and *contextual* faces in which faces were preceded by four non-face object images. Moreover, we investigated how the time components extracted from the FPVS paradigm relate to those extracted from the ERP paradigms. Lastly, we also assessed the relationship between the electrophysiological signals of ERP and FPVS paradigms and the behavioural performance.

### 2.2 Participants

34 participants were tested (4 males, 1 left-handed, mean age: 22.6 ± 2.9, range: 19–30). All participants reported normal or corrected-to-normal vision, and none had reported to have a history of psychiatric or neurological disorders. Observers participated in an EEG session, comprised of two ERP and one FPVS experiments. 29 participants also took part in a behavioural task following the EEG session. Prior to participating in the study, all participants gave their written consent. The study was approved by the local ethics committee and conformed to the Declaration of Helsinki.

One participant was rejected due to excessive noise in the EEG signal resulting in too few trials per condition. The results from 33 participants were retained for the final analysis (22.7 ± 2.9 years, 4 males, 1 left-handed).

### 2.3 Stimuli and procedure

#### 2.3.1 Stimuli and apparatus

Stimuli were displayed at the centre of a VIEWPixx/3D monitor (1920 × 1080-pixel resolution, 120 Hz refresh rate) on a light grey background using the Psychtoolbox in Matlab 2017b (The MathWorks Inc., Natick, MA). Stimuli consisted of photographic images of various man-made objects and images of human faces. The same stimuli were used in previous studies investigating face categorization using the oddball FPVS paradigm (de Heering & Rossion, 2015; Rossion et al., 2015). All objects and faces were presented without isolation from their original backgrounds. While centred within the display, they varied in size, viewpoint, lighting, and background. The stimuli were converted to grayscale, resized to 320 × 320 pixels, and adjusted for pixel luminance and contrast using the SHINE toolbox (Willenbockel et al., 2010). Notably, given that this normalization process is applied to the entire image, the facial features in these natural images retained intentional variations in local luminance, contrast, and power spectrum. The stimuli subtended approximately 6.6 by 6.6 degrees of visual angle.

EEG was acquired by means of BioSemi (Amsterdam, Netherlands) ActiView software with a Biosemi Active-Two amplifier system and 128 Ag-AgCl Active electrodes. Offset was lowered and maintained below 30 mV relative to the common mode sense (CMS) and driven right leg (DRL) by slightly abrading the scalp and adding saline gel. The signal was digitalized at a sampling rate of 1024 Hz. Digital triggers were sent by means of a VPixx Technologies (Saint-Bruno, Canada) screen.

#### 2.3.2 Procedure

A schematic illustration of the ERP and FPVS paradigms as employed is presented in Figure 1. After the implementation of the EEG cap, participants were seated comfortably at a distance of 75 cm from the computer screen. Participants’ head positions were stabilized with a head and chin rest to maintain viewing position and distance constant. Participants were instructed to refrain from any movements during the experiment. Due to the sensitive nature of the ERP signals, participants first completed the ERP experiments, then performed in the FPVS experiment. The order of the *isolated* and *contextual* ERP experiments was randomized for each participant. In all experiments, participants were instructed to fixate on a red cross located on the centre of the screen while continuously monitoring the stimuli presented. In all experiments, participants’ task was to detect brief (500 ms) colour changes (red to blue) of this fixation cross. In the FPVS paradigm, colour changes occurred 10 times within every trial. In the *isolated* ERP paradigm, 24 trials with colour changes occurred randomly among 160 trials with no colour changes. In the *contextual* ERP paradigm, 12 trials with colour changes occurred randomly among 80 trials with no colour changes. This task was orthogonal to the manipulation of interest in the study and was used to ensure that the participants maintained a constant level of attention throughout the experiment.

**Figure 1.**
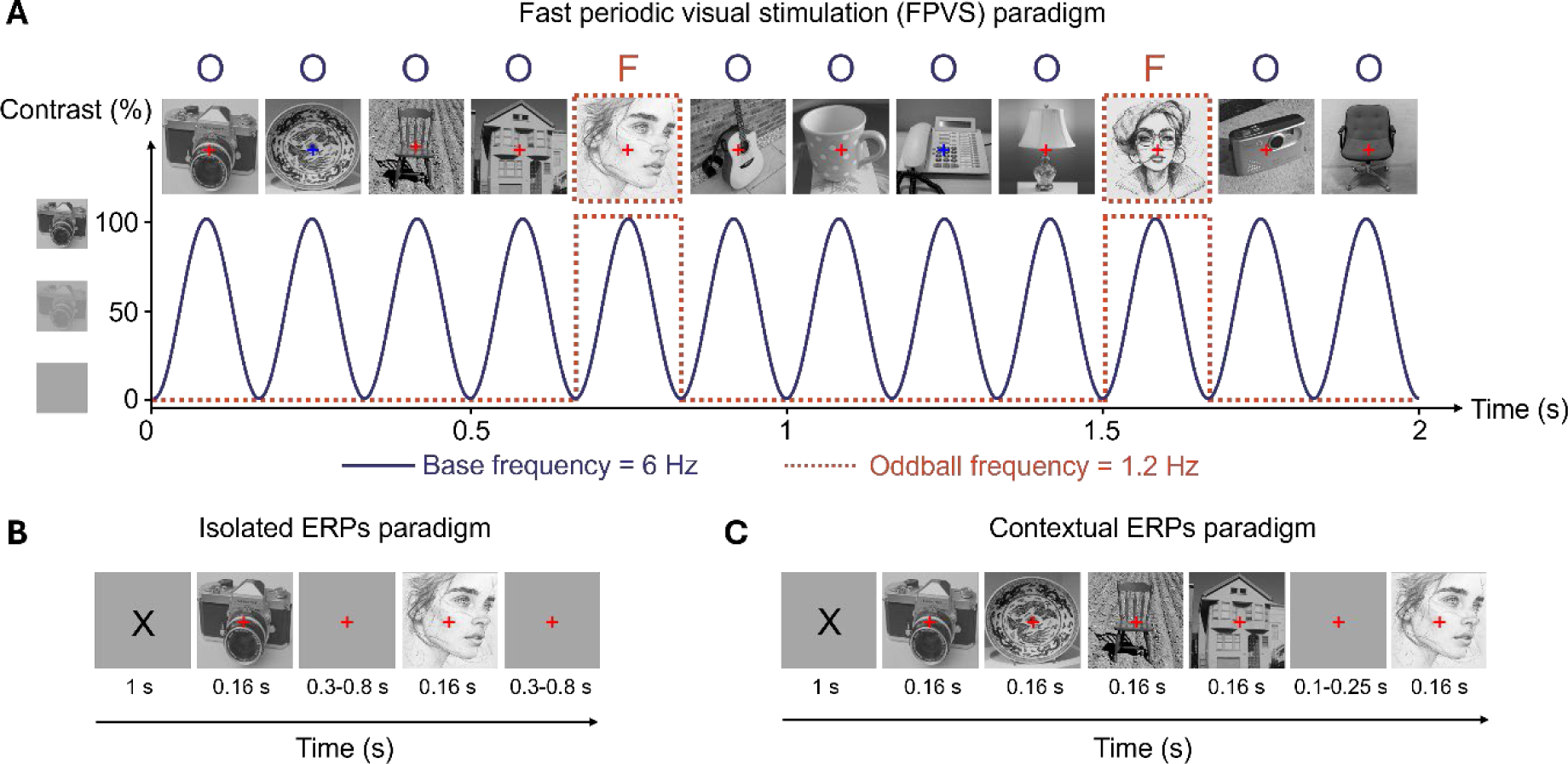
A schematic illustration of the FPVS and ERP paradigms. (A) Natural images of objects were presented at 6 Hz following a sinusoidal contrast modulation. Face images were inserted every 5 stimuli, corresponding to a frequency of 1.2 Hz (= 6 Hz/5). (B) Natural images of objects and faces were presented in isolation. (C) A face image was preceded by four object images. Note that face drawings in the figure differ from real face images used in our experiment due to copyright restrictions of bioRxiv.

In the FPVS paradigm, within each trial, a sequence of visual stimuli was presented through sinusoidal contrast modulation at a base frequency of 6 Hz; hence, each stimulus lasted 0.166 s (i.e., 1000 ms/6). This base stimulation frequency rate was selected because it elicits large periodic brain responses to faces in adults (Alonso-Prieto et al., 2013; Retter et al., 2021). Each trial lasted 68 s, in which the stimuli were presented in sequences lasting 64 s, which were flanked by a 2-s fade-in at the beginning of the sequence, and by a 2-s fade-out at its end. The 2-s buffer time at the beginning and end of each stimulation sequence was used to avoid abrupt onset and offset of the stimuli, which could elicit eye movements. This resulted in a total of 409 images per trial, all of which were different between trials. In each sequence, four non-face objects were presented consecutively, followed by a face, with all stimuli randomly chosen from their respective categories. This arrangement resulted in the oddball faces being presented at a frequency of 6/5 = 1.2 Hz. Consequently, the EEG amplitude specifically at this oddball face stimulation frequency and its harmonics (2.4 Hz, 3.6 Hz, etc.) served as a measure of the brain’s ability to discriminate between faces and objects, as well as its ability to generalize across different facial stimuli. Trials with excessive muscle noise were repeated. Participants completed 4 trials in the FPVS paradigm. During recording the experimenter visually monitored the signal and repeated the trials for which large deflections were observed. During this task subjects allowed to blink whenever it was necessary.

In the *isolated* ERP paradigm, each trial began with a black X mark (≈1° of visual angle) presented at the centre of the screen for 1 s, which was then replaced by a red fixation cross (≈0.3° of visual angle) presented for an interval of random duration between 0.3 – 0.8 s. A face or non-face object was then presented for 0.166 s. The offset of the face or non-face stimulus was followed by an intertrial interval of 1 s. Participants completed 80 trials with face stimuli and 80 trials with object stimuli.

The *contextual* ERP paradigm followed the same procedures as the isolated ERP paradigm except for the following differences. Instead of a single visual stimulus, in each trial a face stimulus, presented for 0.166 s, was preceded by four non-face object images, each presented for 0.166 s. A variable interval duration of 0.1–0.25 s was inserted in between the presentation of face and non-face objects to avoid expectation-related modulation of the neural signal. Participants completed 80 trials.

During both transient ERP paradigms, participants were instructed to refrain from blinking during the presentation of the visual stimuli. During EEG recording the experimenter visually monitored the signal and the participant and repeated trials clearly contaminated by eye-blinks.

A longer version of the Cambridge Face Memory Test (CFMT; Duchaine & Nakayama, 2006) with an added section of 30 difficult trials containing heavily degraded images (CFMT+; Russell et al., 2009) was employed at the end of the EEG session to provide a behavioural measure of facial identity recognition. The test features grayscale cropped male face stimuli with six target and 46 distractor identities. Participants initially are familiarized with images of target identities from diverse viewpoints and subsequently are asked to identify the target image in a three-alternative forced-choice task. As the test progresses, trials become progressively more difficult due to manipulations in illumination, orientation, visual noise, and information availability. We recorded the number of correct responses of each participant across the 102 trials.

### 2.4 FPVS preprocessing and analysis

#### 2.4.1 Preprocessing

All EEG analyses were conducted using Letswave 5 (Mouraux & Iannetti, 2008) and Matlab 2017b (The MathWorks Inc., Natick, MA). Continuous EEG data were first bandpass filtered to exclude frequencies below 0.1 Hz and above 100 Hz using a 4th order Butterworth filter. The signal was then downsampled to 512 Hz. Then, for each observer, 2 × 66 s epochs, which included 2 extra seconds pre- and post-stimulation, were extracted. An independent component analysis using a square mixing matrix algorithm was conducted to remove blink-related noise from each participant’s data (up to two components were selected based on their topography and time-course). Data were then visually inspected and electrodes that presented systematic noise-related deflection over multiple trials were interpolated (with no more than 5% of electrodes interpolated per participant). Subsequently, the data were re-referenced to a common average reference and cropped to an integer number of oddball’s cycles starting 2 s after stimulation onset and ending 2 s before stimulation offset (= 30720 bins).

#### 2.4.2 Frequency domain analysis

The periodic oddball responses were analysed in the frequency domain. Following re-referencing, the amplitude of EEG responses in the frequency–domain was computed using Matlab’s built-in Fast Fourier Transform (FFT) function (N/2 points with normalized amplitudes). To account for baseline variations, baseline-correction was applied to all of the resulting amplitude spectra by subtracting the average of the surrounding 20 bins from each frequency bin, excluding the two immediately neighbouring bins. Signal averaging was performed across 20 (A9-A16, A22-A29, B6-B9) and 10 occipito-temporal electrodes (A10-A12, B7-B11, D31-D32) to encompass channels sensitive to general (base) and face categorization (oddball) responses, respectively. Considering that the periodic response to stimulation is spread over multiple harmonics (i.e. integer multiples of the stimulation frequency), the relevant range of frequency harmonics was determined independently for base and oddball frequencies. Z-scores were calculated after averaging across subjects and electrodes to identify significant harmonics, with significance determined by z-scores exceeding 2.32 (p < .01, one-tailed) for two consecutive harmonics (Z-scores were computed following the same logic as the baseline-correction). Significant responses at the oddball frequency (1.2 Hz) and its harmonics were indicative of face categorization, while responses at the base frequency (6 Hz) reflected a combination of face-related processing and general visual responses. Based on this threshold the oddball response, which indexes implicit neural face categorization, was quantified by summing the first 11 oddball harmonics (i.e., 1.2–13.2 Hz), with the exclusion of the 5th and 10th harmonics due to their overlap with the base stimulation frequency rate (see Figure 2, top panel). The base response remained significant until the 4th harmonic (i.e., 6–24 Hz). As our primary focus was on the face categorization response rather than the overall visual response, we solely assessed the fundamental base frequency (i.e., 6 Hz) to ensure the validity of our experimental manipulation regarding fixated visual input. Topographical maps for oddball and base frequency responses are shown in Figure 3.

**Figure 2.**
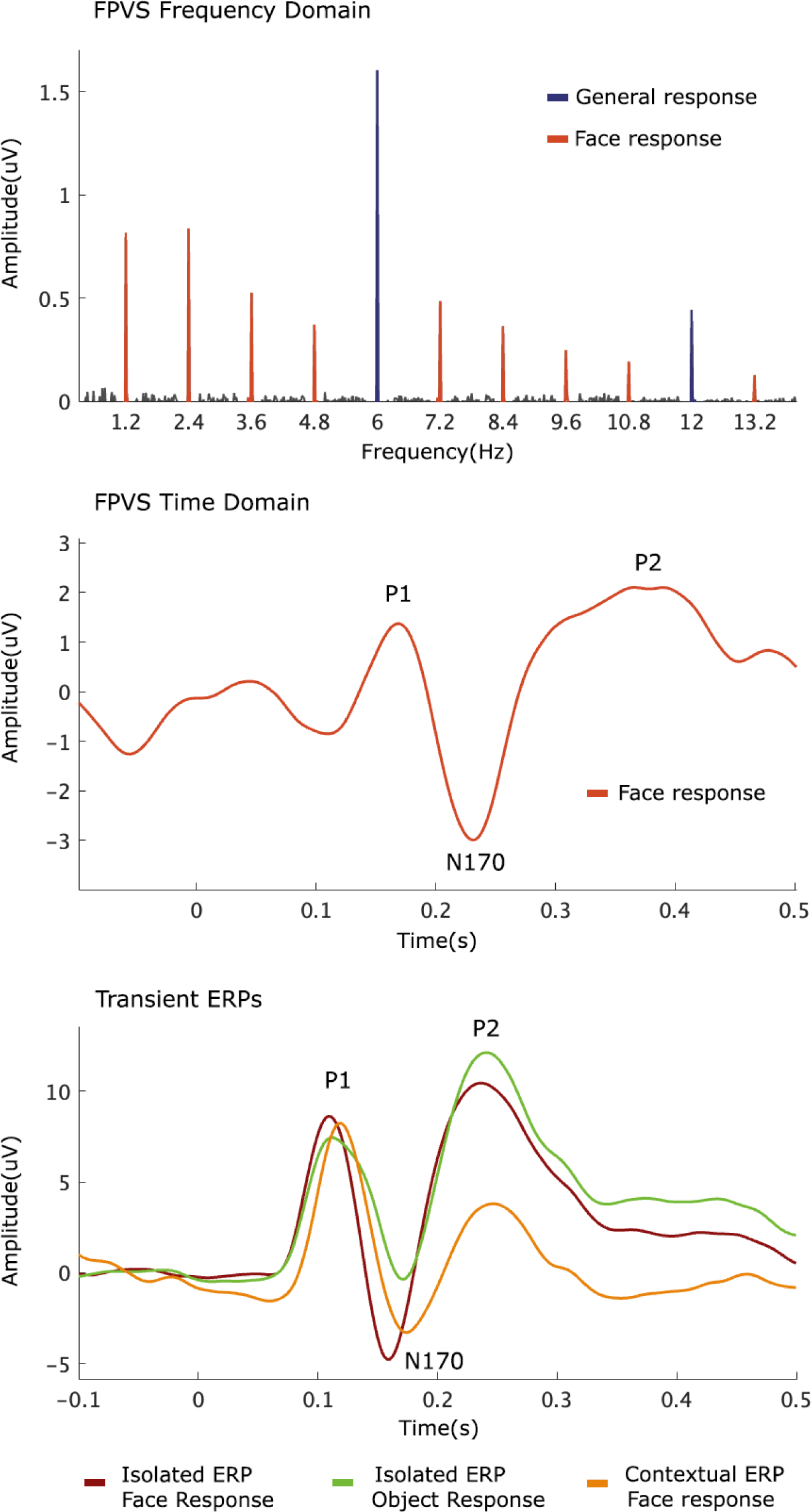
Baseline corrected amplitudes averaged for signals at five right occipito-temporal channels showing significant responses at the oddball stimulation frequency (1.2 Hz) and its harmonics (up to 13.2 Hz), as well as at the base stimulation frequency (6 Hz) and its harmonics (significant up to 24 Hz, but shown until 12 Hz in the figure) (top panel). ERP waveforms averaged for signals at five right occipito-temporal channels showing peaks at P1, N170, and P2 for FPVS time domain (middle panel) and transient ERPs (bottom panel).

**Figure 3.**
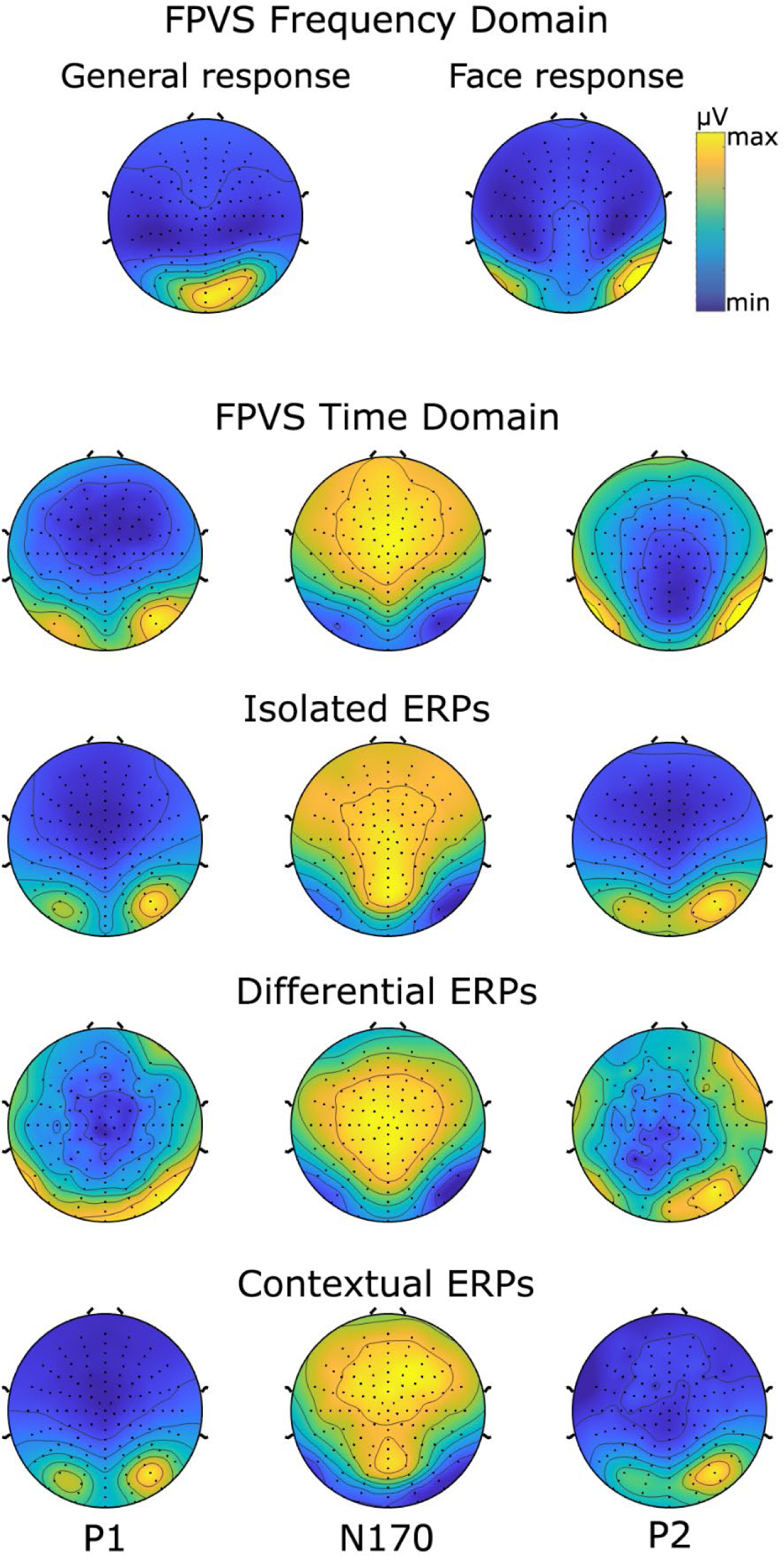
Averaged topographical maps for FPVS frequency domain responses (i.e., general and face-selective), and averaged topographical maps for P1, N170, and P2 peaks in FPVS time domain and transient ERP responses. Averaged topographical map for general and face-selective responses in FPVS frequency domain is based on the sum of corrected amplitudes at significant harmonics.

#### 2.4.3 Time domain analysis

The periodic oddball responses were also analysed in the time domain. The re-referenced EEG data was low-pass filtered with a cutoff frequency of 30 Hz using a fourth-order Butterworth filter. A notch filter (with a width of 0.05 Hz) was then applied to selectively eliminate the influence of the base stimulation frequency and its first five harmonics (ranging from 6 to 30 Hz) from the time-domain waveforms. Subsequently, the filtered data were segmented into smaller epochs lasting 1.67 seconds each, with the initial epochs during the trial’s fade-in period discarded. Baseline correction was performed on these smaller epochs by subtracting the mean response amplitude observed during the 100 ms preceding the presentation of the face image. For each trial, 70 epochs were acquired per participant. These epochs were averaged for each participant and then grand-averaged for display of time-domain data (Figure 2, middle panel).

From these averaged waveforms, ERP components were extracted for each participant at five electrode sites in the right hemisphere, chosen for their known sensitivity to face-selective effects (B7-B11). Consistency in electrode site selection across conditions and individual ERP components was crucial for meaningful regression model comparisons (see statistical analysis section). Visual inspection revealed substantial topographical overlap both amongst the FPVS time domain components; and between the FPVS time domain components and the ERP components (see Figure 3), aligning with prior literature highlighting face-related responses primarily over occipito-temporal sites, with a dominance in the right hemisphere (e.g., Lochy et al., 2019; Rossion, 2014). EEG waveforms revealed three components: a positive component peaking at approximately 160 ms (“P1”), followed by a large negative component peaking at a 220-ms latency (“N170”) and finally by a large positive component that peaks at around 410 ms (“P2”). Despite the delay in components’ latencies (compared to transient ERP components), we will keep referring to these components as P1, N170, and P2 for the sake of simplicity and consistency throughout our manuscript. Finally, the peak-to-peak differences between N170 and preceding P1 (P1-N170) and between N170 and succeeding P2 (N170-P2) were calculated to capture the integrative effects of these components.

### 2.5 ERP preprocessing and analysis

#### 2.5.1 Preprocessing

Continuous EEG data were band-pass filtered between 0.5 Hz and 30 Hz (Butterworth filter, order 4), and downsampled to 512 Hz. Trials with colour changes and button presses were excluded from the analyses. The downsampled data was then segmented into epochs centred on stimulus onset (−2 to 2 s). Electrode coordinates and spline files were assigned for building the 2D scalp maps and the 3D head plots, respectively. Epochs containing blink artifacts within the time window from −0.1 to 0.5 s were removed. Subsequently, EEG data were segmented into 0.6 s epochs (-0.1 to 0.5 s relative to stimulus onset) for each condition (isolated faces, isolated objects, and contextual faces). Baseline correction was performed by subtracting the amplitude in the 0.1 s pre-stimulus period. Noisy channels with systematic deflections exceeding 100 μV over multiple trials were reconstructed using linear interpolation from neighbouring clean channels, with no more than 5% of channels interpolated per participant. The interpolation of occipital channels was maintained consistently across all ERP and FPVS EEG data analysis to prevent biasing of the signal of interest. Epochs containing amplitude exceeding ±75 μV within -0.1-0.5 s time-window were discarded. A common average reference computation was applied to all channels. Finally, averaging was performed across 66 epochs for all subjects, with the subject having the fewest epochs in a specific condition serving as the threshold. Group averaged data of isolated face, isolated object, and contextual ERP waveforms are shown for display of time-domain data (Figure 2, bottom panel).

#### 2.5.2 Time domain analysis

After preprocessing, the EEG waveforms were utilized to extract ERP components for each participant and condition separately (i.e., face and object stimuli conditions in the isolated ERP, face stimulus in the contextual ERP). Differential ERPs were obtained by subtracting the isolated ERPs components to objects from isolated ERPs components to face stimuli. ERPs were extracted within specific time windows corresponding to P1, N170, and P2 at five electrode sites in the right hemisphere (B7-B11), as in the FPVS time domain. Visual inspection of topographies indicated considerable overlap across conditions and individual ERP components (see Figure 3). Peak amplitudes for P1, N170, and P2 were observed approximately between 80 and 140 ms, 150 and 230 ms, and 180 and 250 ms, respectively. These peak amplitudes were then determined and extracted individually for each condition, participant, and electrode of interest. Additionally, as in the FPVS time domain, the peak-to-peak differences between N170 and preceding P1 (i.e., P1-N170) and between N170 and succeeding P2 (i.e., N170-P2) were calculated for all conditions.

### 2.6 Statistical analysis

Statistical analyses were performed in R statistical software (version 3.6.3). Generalized linear models (GLM) with Gamma family and log link function were employed to assess the associations between the amplitude of FPVS oddball frequency responses and the response amplitude from FPVS-time data, isolated ERP, differential ERP, and contextual ERP. Specifically, the FPVS oddball frequency responses were regressed against the P1, N170, P2, P1-N170, and N170-P2 responses separately for each of the four data types (FPVS-time data, isolated ERP, differential ERP, and contextual ERP). This approach yielded a total of 20 models (five predictors by four data types). The models were built using the average amplitude across the 5 right occipito-temporal electrodes.

Bayes factors were used to assess the strength of evidence for the associations between FPVS oddball frequency responses and the responses from FPVS-time, isolated ERP, differential ERP, and contextual ERP analyses. Bayes factors of each model were computed by comparing the predictor model with the intercept-only model (i.e., model with no fixed effect) using the model.comparison() function from the flexplot package (Fife et al., 2021). This comparison helps determine whether including predictors improves model fit and supports any hypothesized association. A Bayes factor greater than 1 suggests evidence in favour of the alternative hypothesis (i.e., the model with a predictor) over the null hypothesis (i.e., the intercept-only model), while a Bayes factor less than 1 indicates evidence in favour of the null hypothesis. Given the necessity to account for multiple comparisons and to mitigate the increased risk of false positives, an evidence boundary of Bayes factor greater than or equal to 10 was chosen to denote significance of the predictor model compared to the intercept-only model (Schönbrodt & Wagenmakers, 2018; Stefan et al., 2019). This higher evidence boundary indicates strong evidence for the presence of an effect, thus reducing the likelihood of erroneously identifying associations. Models passing this significance threshold (BF_10_ ≥ 10) were further contrasted with each other to reveal the hierarchical relationship between them in explaining the FPVS-frequency response. The adjusted deviance-based R-squared values for each model were computed using the adjR2() function from the glmtoolbox package (Vanegas et al., 2023) to assess how effectively the models fit the data. Importantly, the adjusted deviance-based R-squared value can range from negative infinity to 1. A value of 1 indicates a perfect fit, while 0 suggests no improvement over using the response variable. Negative values occur when the model fits worse than no predictors.

To further understand whether the responses elicited by the FPVS paradigm and transient ERP paradigms are equivalent, we also explored the relationship between the ERPs extracted from the time-domain of the FPVS domain, and transient ERPs. The main goal was to compare the responses triggered by these two types of stimulations in the same domain, namely time-domain. To this aim, we used linear models (LM) to regress the FPVS response in the time domain on transient ERPs. This was done for each peak and peak-to-peak separately resulting in 20 models. Bayes factors were computed and the significance threshold (i.e., BF_10_ ≥ 10) was applied as previously. The R-squared values were computed using the model.comparison() function.

Finally, we explored the relationship between participants’ behavioural performance on the CFMT+ and their EEG responses across all paradigms. GLMs (for the FPVS frequency response) and LMs (for the time-domain components) were employed to regress EEG responses onto the accuracy scores of CFMT+. Bayes factors and R-squared values were computed and the significance threshold (i.e., BF_10_ ≥ 10) was applied as previously.

## 3. Results

### 3.1 FPVS frequency response vs. FPVS time domain and transient ERP responses

The models depicting the relationship between the amplitude of FPVS oddball frequency responses and time-domain components across paradigms are shown in Figure 4. Significant associations were observed between FPVS oddball frequency responses and specific time-domain components in each paradigm.

**Figure 4.**
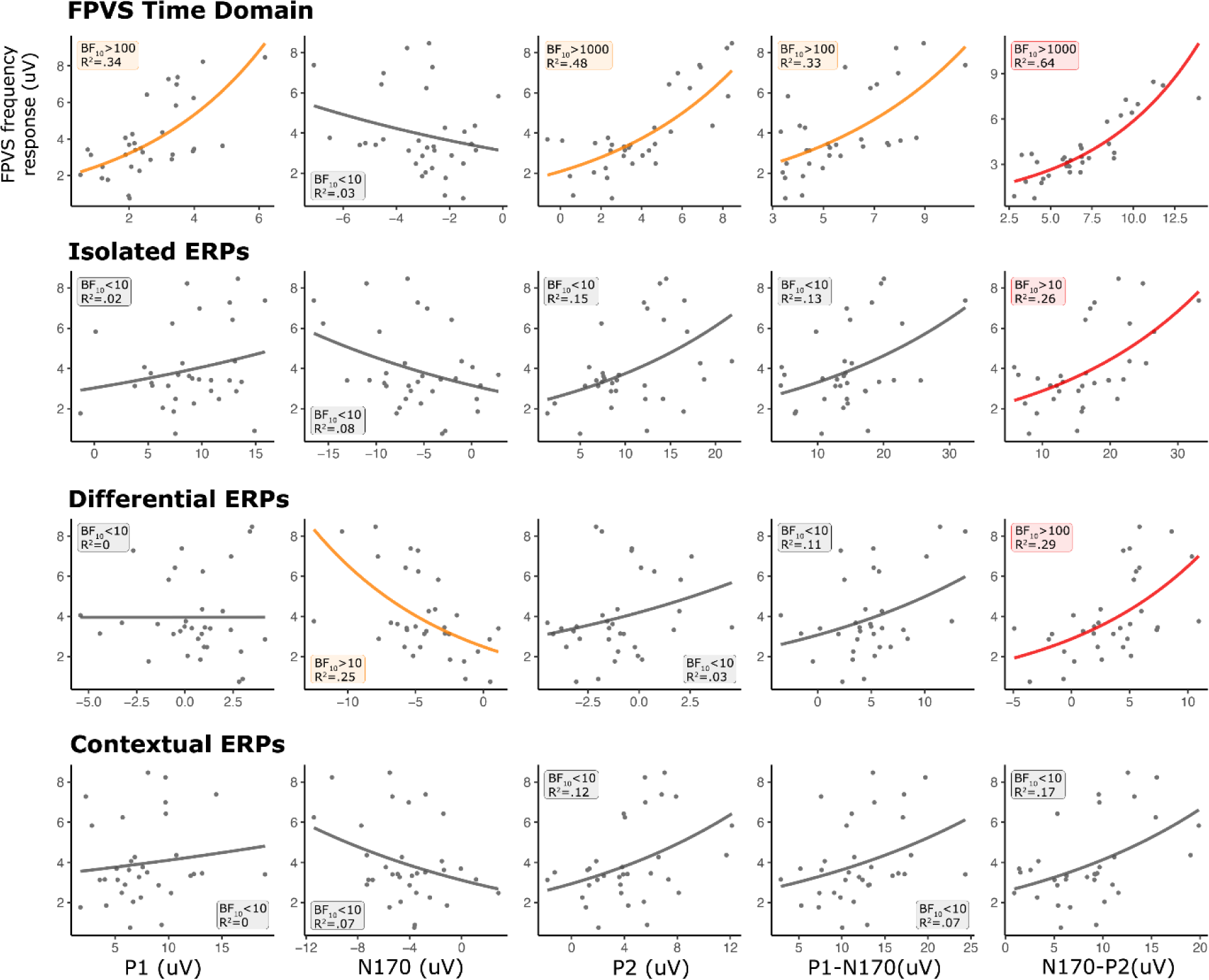
The relationship between the FPVS frequency response and time domain components (P1, N170, P2, P1-N170 and N170-P2) in FPVS time, isolated ERPs, differential ERPs, and contextual ERPs. The individual dots denote individual subjects. The coloured lines denote significance of the predictor model against the intercept-only model. Red lines denote the model with the highest R2 value within a specific paradigm.

Within the FPVS-time components, all components were significantly associated with FPVS frequency response (BF_10_>100, R^2^≥0.33), with the exception of the N170 component (BF_10_<10). Within the isolated ERP components, the only significant association was found between the FPVS frequency response and N170-P2 difference (BF_10_=54.0, R^2^=0.26). Within the differential ERP components, a significant association was found for the N170 component (BF>10, R2=0.25) as well as for the N170-P2 difference (BF_10_>100, R^2^=0.29). Lastly, within the contextual ERP components, no significant association was found, with largest association observed for N170-P2 difference (BF_10_=6.94, R^2^=0.17).

The hierarchical analysis of models provided insights into their respective associations with the FPVS-frequency response. Comparisons between significant models revealed the differences in their associations, shedding light on the relative contributions of different components to the overall variance in FPVS-frequency responses. The N170-P2 in FPVS-time demonstrated the strongest association, followed by the P2 in FPVS-time, while the N170-P2 in isolated and differential ERP, N170 in differential ERP, the P1 and the P1-N170 in FPVS time showed comparable associations. This hierarchical delineation underscores the differential impact of various components on explaining the variance in FPVS-frequency responses, with the N170-P2 difference in FPVS-time emerging as the most influential predictor.

### 3.2 FPVS time domain vs. transient ERP responses

The models depicting the relationship between the components extracted from the time-domain of the FPVS response and those obtained from transient ERP are shown in Figure 5. Within the isolated ERP condition, significant associations were found for the P2 (BF_10_>10), P1-N170 (BF_10_>100), and N170-P2 (BF_10_>1000). Within the differential ERP condition, a significant association was observed for the P1-N170 difference (BF_10_>10). Finally, within the contextual ERP condition, a significant relationship was found for the P2 component (BF_10_>100) and for the N170-P2 different (BF_10_>10). In line with results above, this pattern underscores the significance of the N170-P2 difference, which emerges as a consistent predictor across paradigms (except in the differential ERP condition).

**Figure 5.**
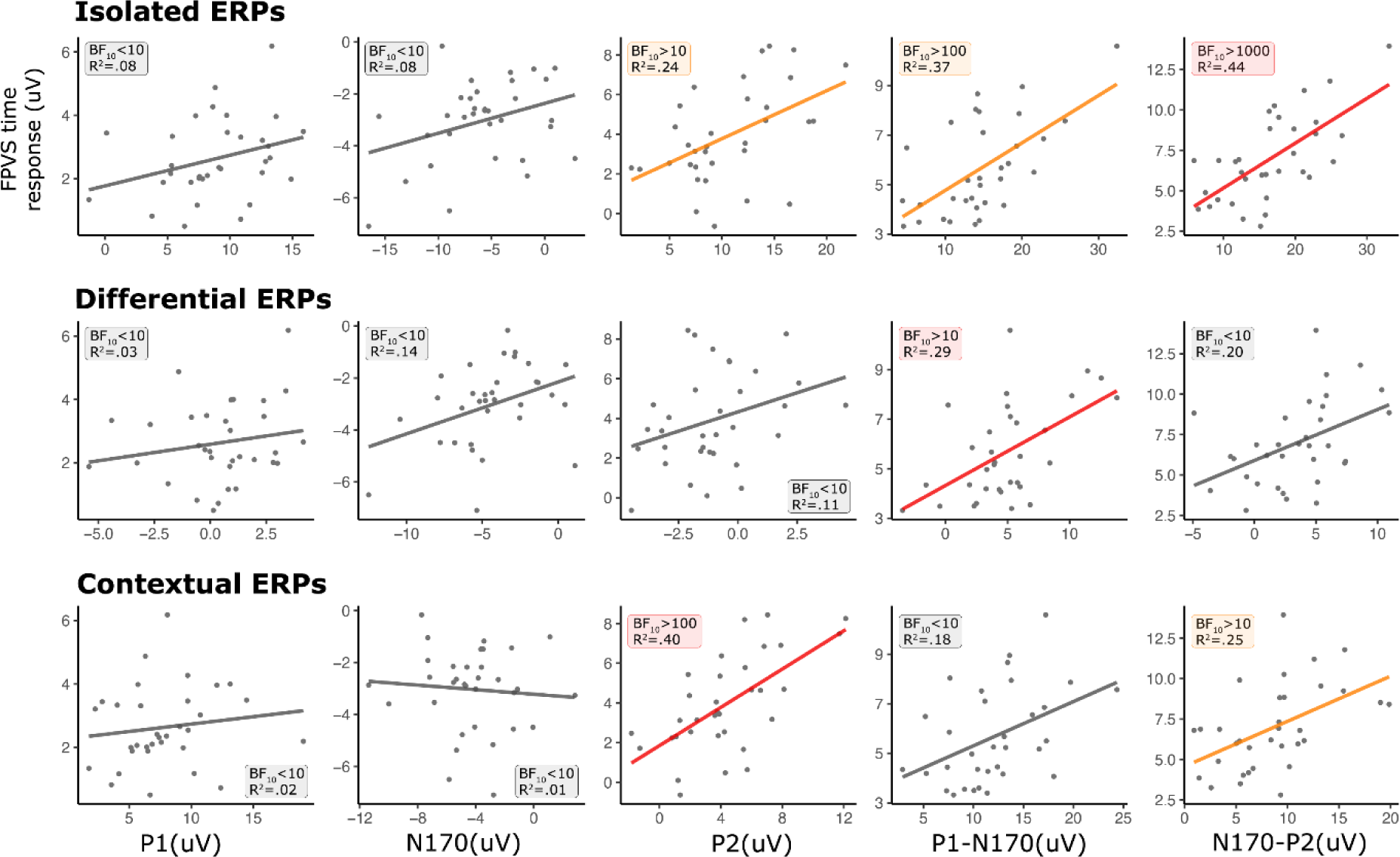
The relationship between the FPVS time domain components and ERP time domain components. The individual dots denote individual subjects. The coloured lines denote significance of the model against the intercept-only model. Red lines denote the model with the highest R2 value within a specific ERP condition.

### 3.3 Neural face categorisation responses vs. behavioural performance

Finally, the regression models assessing the relationship between the accuracy scores in the CFMT+ revealed that the accuracy in the CFMT+ was not significantly associated with any of the EEG responses including the frequency response and time domain responses.

## 4. Discussion

In the current study we aimed to assess whether the face categorization response obtained through the FPVS paradigm was related to the face sensitive N170 ERP component or was associated with different neural responses. To this aim, we assessed the relationship between the FPVS response in the frequency domain and the ERP components extracted from the FPVS response in the time domain as well as those elicited by *isolated* faces, those obtained by removing object responses from face responses (i.e., *differential* face response) and those elicited by faces presented after a short series of objects (i.e., *contextual* face response). For each time-domain measure we extracted the P1, N170 and P2 peaks, as well as the P1-N170 and N170-P2 peak-to-peak responses. As the neural responses were much more prominent in the right hemisphere than the left hemisphere, and that face processing it typically right lateralized (e.g., for a review see: Rossion & Lochy, 2022), we focused our analysis on a right occipito-temporal region of interest (ROI). Overall, our data show that the N170 ERP component holds little to no explanatory power over the FPVS frequency response, which was most consistently and strongly explained by the peak-to-peak difference between the N170 and the P2 components.

### 4.1 FPVS frequency domain vs. FPVS time domain

When analysing ERP components from the FPVS time domain, we found a significant relationship with the frequency response for all responses, except for the N170 component, while the N170-P2 difference showed the largest explanatory power. Within the context of isolated face responses, only the N170-P2 difference showed a significant relationship with the FPVS frequency response. In the case of differential face responses, although the N170 was significantly linked to the frequency response, this association was weaker than that observed with the N170-P2 difference. Lastly, regarding contextual face responses, none of the components displayed a significant relationship with the frequency response.

The observation that the FPVS frequency response correlates more strongly with the N170-P2 response compared to other responses may be attributed to the method used to obtain the frequency spectrum, namely the fast Fourier transform (FFT). This procedure assesses the presence of sinusoidal waves of different frequencies within a given signal. The amplitude of a wave is determined by the difference between the lowest point in the trough and the highest point in the peak. Consequently, a peak-to-peak difference would likely have a greater impact on the frequency amplitude compared to a single peak. However, the reasons why the P1 and P2 single peaks extracted from the FPVS time-domain, but not the N170, were significantly related to the frequency response warrants further explanation.

One possible explanation is that within the context of FPVS, the rapidity of the neural response to face stimuli might have varied significantly among different face exemplars and different observers. We presented six stimuli per second using natural images, where faces and non-face objects were embedded in diverse backgrounds. Moreover, faces had different head orientations, expressions, age, or gender. This variability, along with potential factors such as occasional lapses in attention, participants blinking coinciding with face presentation, and general fatigue, might have made some face-stimuli within the sequence more difficult to detect. As shown by Latinus and Taylor (2006), difficulties in face processing can modulate ERP components’ latency (see also Jemel et al., 2003; Németh et al., 2014). Their findings indicate that detecting a face in a Mooney-face, which poses a greater challenge for detection due to their simplified nature and lack of detailed features compared to photographic faces, caused a greater delay in the N170 compared to the P1 and P2 components (Latinus & Taylor, 2006). Greater latency variability leads to less periodic occurrence of a component, making it less conspicuous in the frequency domain.

A recent FPVS study by Retter and colleagues (2020) has shown that when observers fail to detect a face amongst other non-face objects, the FPVS time-domain response obtained lacks any discernible ERP component. However, there is currently no information available regarding instances where a face is detected with difficulties, which might be indexed by longer latencies or higher uncertainty. Further research is needed to explore the role of stimulus-driven difficulties on the FPVS response in both the time and frequency domain. Additionally, when we extract the N170 from the time domain, we average the responses following each face presentation, which can result in a less precise final peak due to latency variations. Therefore, it is reasonable to anticipate that the correlation analysis that we carried out had more contamination by noise from the time domain, and less contribution of the N170 in the frequency domain. This avenue should be explored by future studies, for example by using slower presentation frequencies or segregated images of faces and non-face objects. Such approaches could potentially reduce the difficulties of neural face categorization. Clarifying this aspect is important for a better understanding of the FPVS response. If confirmed, it could suggest that the frequency response across different observers may reflect different stages of face categorization. While some subjects could conform to our group-level observation, others may have responses driven more strongly by the N170 component.

### 4.2 FPVS frequency domain vs. transient ERP responses

When comparing the FPVS frequency response to face responses elicited in ERP paradigms, we found that for both the differential and isolated face response, the N170-P2 difference showed the highest explanatory power. Along with the results from FPVS-time domain, this suggests that the frequency response is an integrated measure of early and later face-related responses.

On the other hand, no ERP component from the contextual face response showed a significant relationship with the FPVS frequency response. We implemented the contextual ERP paradigm to mimic the FPVS presentation conditions, where a face-stimulus is preceded by a short sequence of non-face stimuli. We reasoned that ERPs in this case would more closely relate to the FPVS frequency response as in both cases a face-stimuli is presented within context. We hypothesise that a lack of relationship might be attributed to the methodological choice of a short and variable inter-stimulus interval (i.e., ISI) between the last object and the face-stimulus. These parameters were chosen to reduce the predictability of the face onset, and, at the same time, to minimize the delay between the two types of stimuli. However, this might have caused other issues. Notably, the short ISI might have caused contamination of the face response by the object response. Importantly, as the ISI varied across trials, it is possible that this contamination might have led to inconsistent components onset within the face response. This in turn could have introduced an amount of noise that overpowered the signal. In line with this reasoning is the observation that the amplitude of the N170 and P2 ERP components in the contextual condition is significantly weaker than in the isolated condition. If the components onset was unstable across repetitions, the average peak might be spread over a wider time-window, and, in turn, be weaker. Consequently, the weak face signal did not correlate well with the highly face-sensitive FPVS frequency response.

In terms of the N170 component, we found a significant relationship only for the one extracted from the differential face response. We suggest that the significant relationship observed with the N170 component extracted from the differential face response is because this ERP response aligns more closely with what the FPVS frequency response reflects: a differential response free from contamination related to non-face objects.

### 4.3 FPVS time domain vs. transient ERP responses

To attempt to better understand the relationship between the FPVS response and the differential N170, we regressed the ERP components extracted from the different ERP paradigms on those obtained from the FPVS time-domain data. We found that the FPVS-N170 was not related to any of the N170 component obtained through ERP paradigms, not even the differential one.

When the same analysis was performed also for P1, P2 and the two peak-to-peak differences, we found that, as the N170, also the FPVS-P1 was not related to any of the P1 extracted from the responses to ERP paradigms. FPVS-P2 and N170-P2 were both significantly related to those from isolated and contextual ERPs. On the other hand, the P1-N170 differences recorded in FPVS was related to the one obtained from contextual and differential ERPs. The lack of a relationship for the two early components, P1 and N170 might suggest that the procedural nature of the FPVS paradigm leads to an alteration of the information carried by earlier components, as they are the ones closer in time to the response to the preceding object.

In one of their original papers on frequency tagging in the context of face processing, Jacques et al. (2016) noted that although they look similar in topography and latency, it cannot be assumed that the components extracted from the temporal domain of the FPVS responses are comparable to those obtained from more traditional ERP paradigms. It was argued that the deflections observed in the FPVS time domain represent an inherent contrast response between faces and non-face stimuli (i.e., a face-selective response), rather than an event-related potential (ERP) component in response to the sudden appearance of a face against a uniform background. Our results support this idea only for the earlier components such as P1 and N170, while the peak-to-peak differences and the P2 appear to be more similar across measures.

### 4.4 Neural face categorisation responses vs. behavioural performance

A question that remains unanswered is how the FPVS face categorization response related to behavioral performance. In a recent study, Retter et al. (2020) observed that at the group level, conditions leading to higher face detection performance are those leading to a stronger FPVS frequency response. Nevertheless, whether the neural response could predict behavioural performance at the single subject level remains unknown. Relating this measure and neural responses obtained through ERP paradigms to different behavioral test assessing different stages of human face categorization might shed some light on the result of the current work. In our study, we solely utilized a face recognition task, correlating it with both the FPVS frequency response and various ERP components. Our findings revealed no significant relationship between the FPVS frequency response and different ERP components with behavioral performance in this task. However, it is important to note that our investigation was limited to a single behavioral task. Other behavioral tasks that more closely align with the cognitive processes captured by the FPVS and ERP paradigms may yield stronger correlations with neural responses. Exploring these alternative behavioral measures could provide further insights into the relationship between neural activity and behavioral performance in face categorization.

### 4.5 Right vs left hemisphere in face processing

Consistent with a large body of the face processing literature, our analysis of face categorization responses was mainly focused on the EEG signals coming from the right hemisphere. While both hemispheres contribute to face processing, research suggests that the right hemisphere generally exhibits greater dominance in this domain (Barton, 2008; Cohen et al., 2019; Rossion & Lochy, 2022). Neuroimaging studies have revealed that face-selective areas in the fusiform gyri tend to be more pronounced in the right hemisphere than in the left (Kanwisher et al., 1997; McCarthy et al., 1997). Lesion studies also support the dominance of the right hemisphere, as damage to this hemisphere is often necessary and can be sufficient to induce deficits in facial recognition and emotion processing (e.g., Barton, 2008; Bouvier & Engel, 2006; Cohen et al., 2019; De Renzi et al., 1991; Landis et al., 1986; Sergent & Signoret, 1992). Similarly, a substantial number of EEG studies on face processing based the interpretation of their findings on the right hemisphere as the response was stronger and more reliable in the right than in the left (e.g., Bentin et al., 1996; Lochy et al., 2019; Nguyen & Cunnington, 2014; Rossion & Jacques, 2011; Rousselet et al., 2008, 2014). Consistent with the existing literature, we found stronger peak and trough amplitudes in the right than in the left hemisphere in our face categorization task across paradigms and reported our main results on the right hemisphere. As the focus of our study was to uncover whether the face-specific FPVS frequency response could be a proxy for the isolated time domain components such as the face-sensitive N170, the further investigation into left-right differences in face categorization was out of scope of this research.

## 5. Conclusions

Our study pioneers the investigation into the neural mechanisms indexed through the FPVS frequency response, a critical inquiry for researchers adopting this approach to study neural face categorization. Our results challenge the assumption of a straightforward relationship between rapid neural face categorization, as captured by the FPVS paradigm, and the face-sensitivity indexed by the N170 component. We provide empirical evidence refuting the notion that the FPVS frequency response should serve as a proxy for the N170, suggesting instead that it likely represents a later stage in the face processing continuum. Further exploration is necessary to fully elucidate this relationship, such as studies examining how the FPVS frequency response behaves under specific experimental manipulations compared to known ERP components. Despite its promise, the FPVS technique capture complex neural responses that requires thorough investigation to enable researchers to fully harness their mechanistic and theoretical interpretation. Future studies are also necessary to investigate whether task constraints (e.g., face discrimination) would modulate the relationship we observed between FPVS frequency and face sensitive N170 responses.

Our study aimed to enhance understanding of the FPVS response in the context of face categorization by directly comparing frequency to time domain responses. Our findings underscore the N170-P2 difference as the primary predictor of the FPVS frequency response, contrasting with the limited correlation observed between the FPVS frequency response and the N170 component. Although the complexity of the FPVS frequency response and the spatial resolution limitations of EEG pose challenges to theoretical interpretation, our methodological findings invite to caution against potential (unfounded) theoretical claims that the FPVS frequency response is solely rooted in the N170 component.

## 6. Data and Code Availability

The pre-processed EEG data and behavioural data that support the findings of this study are available on request from the corresponding author.

## 7. Author Contributions

FZKY: Conceptualisation, Methodology, Formal analysis, Investigation, Data Curation, Writing—Original Draft, Writing—Review & Editing, and Visualisation. LS: Conceptualisation, Methodology, Software, Formal analysis, Investigation, Data Curation, Writing—Review & Editing, and Visualisation. RC: Conceptualisation, Methodology, Writing—Review & Editing, Funding acquisition, Resources, and Supervision.

## 8. Funding

This study was supported by the grant no 10001C_201145 from the Swiss National Foundation awarded to RC.

## 9. Declaration of Competing Interests

The authors declare no competing interests.

## Notes

### Competing Interest Statement

The authors have declared no competing interest.

